# Genomic epidemiology of *Plasmodium knowlesi* reveals putative genetic drivers of adaptation in Malaysia

**DOI:** 10.1101/2024.04.10.588982

**Authors:** Jacob A F Westaway, Ernest Diez Benavente, Sarah Auburn, Michal Kucharski, Nicolas Aranciaga, Sourav Nayak, Timothy William, Giri S Rajahram, Kim A Piera, Kamil Braima, Angelica F Tan, Danshy Alaza, Bridget E Barber, Chris Drakeley, Roberto Amato, Edwin Sutanto, Hidayat Trimarsanto, Nicholas M Anstey, Zbynek Bozdech, Matthew Field, Matthew J Grigg

**Author notes:** Equal contribution (Matthew Field, Matthew J Grigg). Author contributions Conceptualisation and methodology: MG, MF, ZB, SA, NMA, EB, RA, and JW. Project administration: MG, MF, ZB, SA, NMA, and JW. Supervision: MG, MF, ZB, EB, NMA, and SA. Funding acquisition: MG, MF, ZB, SA, NMA, BB, CD, GR, and TW. Data curation: DA, JW, MG, and SN. Formal analysis, visualisation, and software: JW, EB, MF, ES, HT, MG, and SA. Resources: MG, ZB, TW, GR, BB, and NMA. Investigation: KP, MK, NA, SN, AT, and KB. Writing – original draft: JW. Writing – review and editing: MG, MF, ZB, SA, NMA, EB, KB, MK, NA, ES, HT, CD, and JW.

## Abstract

Sabah, Malaysia, has amongst the highest burden of human *Plasmodium knowlesi* infection in the country, associated with increasing encroachment on the parasite’s macaque host habitat. However, the genomic make-up of *P. knowlesi* in Sabah was previously poorly understood. To inform on local patterns of transmission and putative adaptive drivers, we conduct population-level genetic analyses of *P. knowlesi* human infections using 52 new whole genomes from Sabah, Malaysia, in combination with publicly available data. We identify the emergence of distinct geographical subpopulations within the macaque-associated clusters using IBD-based connectivity analysis. Secondly, we report on introgression events between the clusters, which may be linked to differentiation of the subpopulations, and that overlap genes critical for survival in human and mosquito hosts. Using village-level locations from *P. knowlesi* infections, we also identify associations between several introgressed regions and both intact forest perimeter-area ratio and mosquito vector habitat suitability. Our findings provide further evidence of the complex role of changing ecosystems and sympatric macaque hosts in Malaysia driving distinct genetic changes seen in *P. knowlesi* populations. Future expanded analyses of evolving *P. knowlesi* genetics and environmental drivers of transmission will be important to guide public health surveillance and control strategies.

**Author Summary:** The zoonotic *P. knowlesi* parasite is an emerging, yet understudied, cause of malaria in Southeast Asia. Sabah, Malaysia, has amongst the highest burden of human P. knowlesi infection in the country, however, the region is currently understudied. Thus, we produced a collection of high-quality *P. knowlesi* genomes from Sabah, and in combination with publicly available data, performed an extensive population genetics analysis. Our work contributes novel insights for *Plasmodium knowlesi* population genetics and genetic epidemiology.

## Introduction

Zoonotic transmission of the macaque parasite *Plasmodium knowlesi* has emerged as the most common cause of human malaria in Malaysia and parts of western Indonesia (1, 2, 3). *P. knowlesi* infections can cause severe, life-threatening malaria, with a case fatality similar to that of *P. falciparum* in Southeast Asia despite comparatively lower levels of parasitemia (4, 5). The recent increased reporting of *P. knowlesi* infections in Southeast Asia has been strongly linked with the encroachment of humans on previously intact habitats of their natural macaque reservoir hosts (6). Zoonotic transmission of *P. knowlesi* is thought to occur largely in response to increasingly fragmented landscapes as a result of land clearing and associated agricultural activities, with increased exposure in at-risk workers and local populations in endemic areas to both pig-tailed (*Macaca nemestrina)* and long-tailed (*M. fascicularis)* macaques, and the *Anopheles* Leucosphyrus Group mosquito vectors (7, 8). Worryingly, in contrast to other human *Plasmodium* species, national WHO malaria elimination goals in Southeast Asia are threatened by the inability of public health measures to target macaque host reservoirs for *P. knowlesi* (2). Furthermore, conventional prevention measures such as insecticide-treated bed nets used successfully for other *Plasmodium* Spp in the region are limited for *P. knowlesi* zoonotic infections, primarily acquired at the forest-edge during agricultural work activities (9, 10).

Insights gained from genomic analyses of human malaria parasites have advanced our understanding of basic disease biology, drug resistance and malaria epidemiology (11). Large-scale, collaborative efforts to produce publicly available population-level whole genome data for *Plasmodium* species of interest, have produced over 20,000 *P. falciparum* (12) and ∼1,800 *P. vivax* (13) genomes. In contrast, *P. knowlesi* currently has fewer than 200 whole genomes available from a limited geographic distribution (14, 15, 16, 17, 18). Only 16 reported *P. knowlesi* genomes are described from the state of Sabah in East Malaysia, despite this area representing among the highest reported number of *P. knowlesi* cases and disease burden globally to date (19).

Previous studies of *P. knowlesi* population genetics in Malaysia have identified three genetically divergent populations using a combination of whole-genome sequencing (20) and microsatellite genotyping (21). One of these populations is restricted to Peninsular Malaysia, whilst the other two are found in Malaysian Borneo. The two overlapping clusters in Malaysian Borneo are derived from the separate macaque reservoir hosts: from long-tailed macaques (*Macaca fascicularis* [*Mf*]) and pig-tailed macaques (*M. nemestrina* [*Mn*]) (22). We refer to these clusters as *Mf* (cluster 1), *Mn* (cluster 2) and *Peninsular* (cluster 3) throughout this manuscript. Despite these clearly-defined, genetically divergent populations, previous work further identified distinct subpopulations within the different clusters (15), with evidence of recent positive directional selection (20) and large genetic introgression events between the subpopulations linked to mosquito vectors (15). In this context, introgression refers to the transfer of genetic information from one cluster to another, resulting from hybridisation and repeated backcrossing. This evidence suggests that *P. knowlesi* population structure is changing, with changes hypothesised to occur as a result of rapidly altering forest and agricultural ecosystems in Malaysia.

To address this gap in our understanding of the genomic make-up *P. knowlesi* and patterns of population structure across Malaysia, we performed whole genome sequencing on 94 new human infections from diverse landscapes across Sabah, East Malaysia (19). The newly produced data were combined with 108 publicly available *P. knowlesi* genomes derived from clinical infections across Malaysia (14, 20). Leveraging the additional isolates from Sabah, our objective was to perform a comprehensive evaluation of *P. knowlesi* population structure with a dataset that better represents the distribution of symptomatic infections from passive case detection across Malaysia. We combined genomic data with environmental land cover classification data surrounding knowlesi malaria case villages to better explore the relationship between the genomic and ecological features in Sabah associated with the transmission of *P. knowlesi* populations. These integrated analyses aim to provide insights to assist in the development of future public health interventions and genomic surveillance efforts.

## Results

The 94 newly sequenced *P. knowlesi* whole genomes all originated from the state of Sabah, encompassing human infections from 11 administrative districts, including 22 infections from Kota Marudu and 14 from Kudat (Supplementary Table 1). These genomes had an average number of reads per sample of 66,415,402.53, with 30.27% mapping to the PKA1-H.1 reference genome (23). The average sequencing depth of these new genomes was 84.4, with 82.9% of the bases in the reference genome covered. The 108 high-quality publicly available *P. knowlesi* genomes from NCBI were derived predominantly from human infections (with six laboratory strains passaged through macaques) across different districts from both Peninsular Malaysia (n=33) and East Malaysia, including the geographically distinct neighbouring eastern state of Sarawak (n=59) in addition to a small number from Sabah (n=16) in Malaysian Borneo. After filtering for clonality, missingness and minor allele frequency, 52 of the newly sequenced genomes remained, and another 100 genomes from the publicly available data. The combined 152 *P. knowlesi* genomes included in the population-based analyses consisted of: 54 from Sabah, 66 from Sarawak, and 32 from Peninsular Malaysia. Joint genotyping initially identified 1,542,627 single nucleotide polymorphisms (SNPs), which after filtering resulted in 357,379 SNPs taken forward for downstream analyses.

### High prevalence of polyclonal *P. knowlesi* infections in Malaysian Borneo, but infrequent superinfection

Given that *P. knowlesi* parasites are haploid in the blood stage of host infection, the presence of multiple alleles at given loci is indicative of a multiple clone (polyclonal) infection. The within-isolate fixation index (*F*_WS_) was used to measure the genetic complexity within infections. The *F*_WS_ score ranges from 0 to 1, with increasing values reflecting increasing clonality (24). At a commonly applied threshold of *F*_WS_ < 0.95, 13.4% (n=27/201) of infections were polyclonal. The highest proportion of polyclonal infections was observed in Sarawak (17.6%, n=13/74), followed by Sabah (10.6%, n=10/94), and Peninsular Malaysia (12.1%, n=4/33). There were no statistically significant differences demonstrated for the mean *F_WS_* between the three states (*Figure 1*.A).

**Figure 1.**
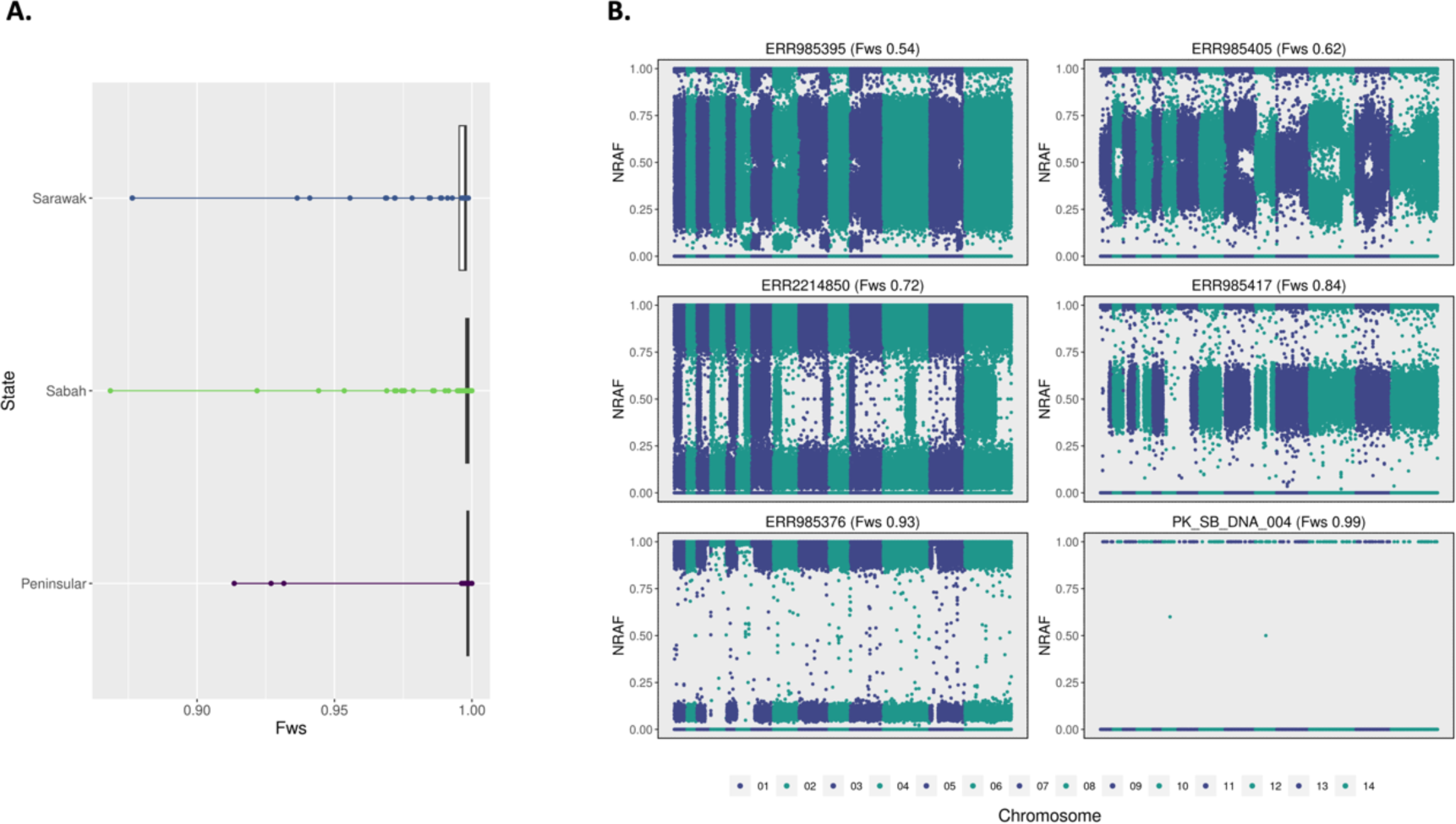
Comparable within-isolate genetic complexity across geographic regions. A: Boxplots depicting the distribution of within-infection diversity (*F_WS_*) across three regions (*Sabah*, *Sarawak*, and *Peninsular*). B: Manhattan plots of the within-isolate non-reference allele frequencies (NRAF) across the genome for six Malaysian *P. knowlesi* infections ranging from low diversity (*F_WS_*=0.99) to high diversity (*F_WS_*=0.54) and with varying levels of within-host relatedness.

Closer inspection of the relatedness between the clones within the polyclonal infections was undertaken using within-isolate, non-reference allele frequency (NRAF) plots. As illustrated in the examples in *Figure 1.B*, a variety of relatedness patterns were observed, including highly distinct clones (e.g. ERR985376 [*F*_WS_ = 0.93]) which, considering obligate recombination of strains in the mosquito host, likely represent superinfections (multiple mosquito inoculations). Other polyclonal infections comprised more related clones (e.g. ERR985395 [F_ws_ = 0.54]) that likely reflected co-transmission of siblings (single mosquito inoculation). Given adequate genetic diversity in a population (not inbred), superinfections with highly related clones are unlikely. Amongst the complex infections, 18.5% (n=5/27) appear to be superinfections. Three of the superinfections were from Malaysian Borneo, including Sabah (Pitas), and Sarawak (Kapit and Betong). The remaining two superinfections were from Peninsular Malaysia, (Temerloh and Kuala Lipis districts). All superinfections appeared to have two clones present.

### *P. knowlesi* new genomes from Sabah belong predominantly within the *Mf* cluster

Previous genetic studies have described distinct genetic clustering of *P. knowlesi* into a geographic Peninsular-Malaysia sub-population, and two Malaysian-Borneo macaque-associated subpopulations; *M. fascicularis* (*Mf*) and *M. nemestrina* (*Mn*) (17, 22, 25). We sought to determine the genetic clustering patterns of the Sabah genomes relative to infections from Sarawak and Peninsular Malaysian. Neighbour-joining analysis based on identity by state (IBS) was undertaken on the 152 low complexity *P. knowlesi* genomes from across Malaysia, revealing three clusters (*Figure 2.A*). The newly sequenced *P. knowlesi* samples originating from Sabah group predominantly within the *Mf* cluster (82.3%, n=43/52); the remaining (17.3%, n=9/52) infections clustered within the *Mn* clade, similar to the proportions of the 16 samples from Sabah previously described by Divis et al. 2017 (*Mf* = 86.6%, *Mn* = 13.4%) (21). ADMIXTURE analysis revealed greatest likelihood of 3 sub-populations amongst the 152 infections, confirming the patterns observed with neighbour-joining analysis (*Figure 2.C*).

**Figure 2.**
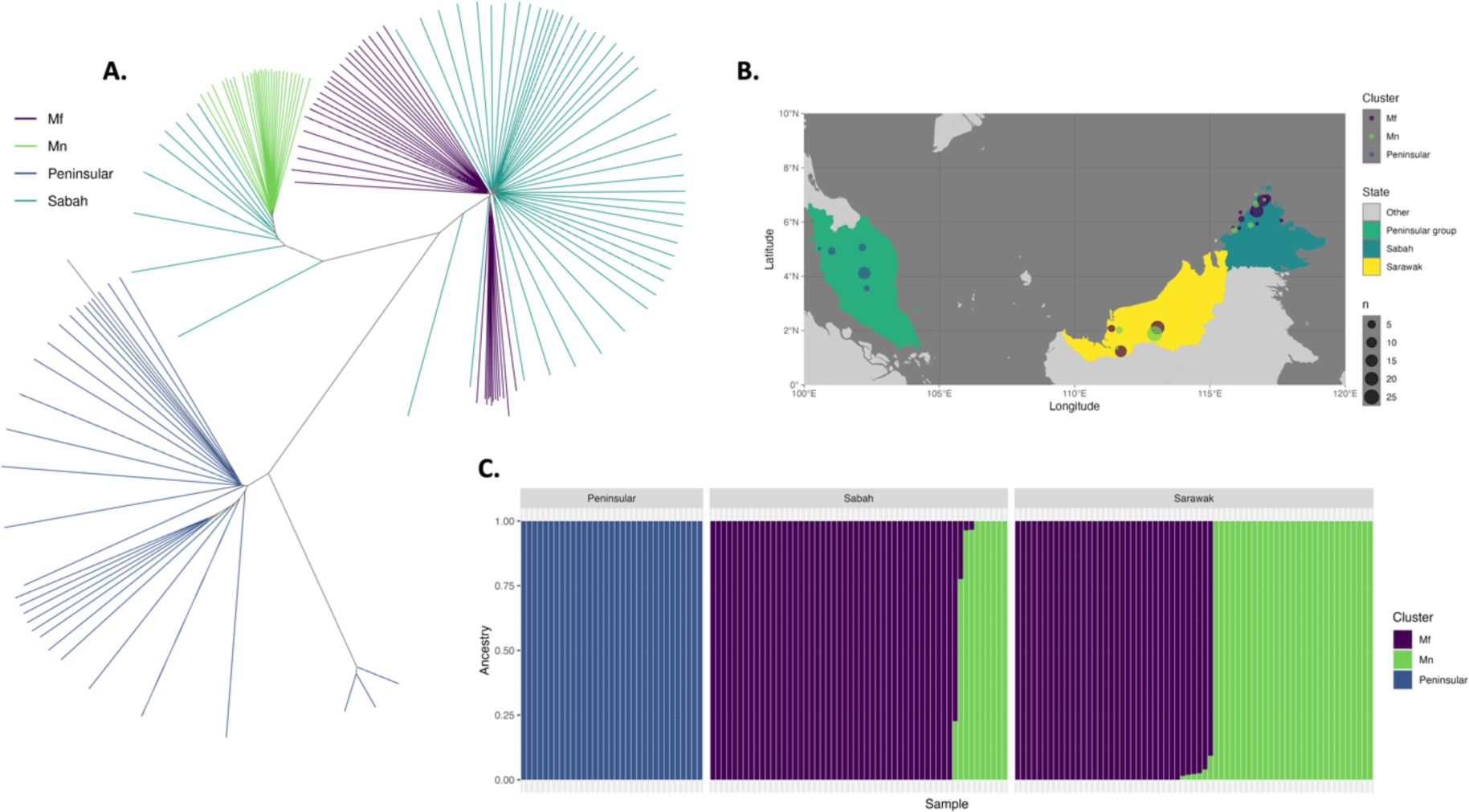
Geographic overlap and evidence of shared ancestry between the *Mf* and *Mn P. knowlesi* clusters. A: Unrooted neighbour-joining tree based on identity by state (IBS) depicting three predominant genomic clusters of *P. knowlesi* across Malaysia, specifically the Peninsular Malaysia sub-population (*Peninsular*), and Malaysian-Borneo macaque-associated subpopulations of *Macaca fascicularis* (*Mf*) and M. *nemestrina* (*Mn*). The new isolates from Sabah are labelled separately (*Sabah*). B: Map of Malaysia showing the geographic distribution and number of samples, and genomic clusters across Malaysian-Borneo (right) and Peninsular-Malaysia (left). C: Bar plot illustrating the proportionate ancestry to each of 3 (K) subpopulations determined by ADMIXTURE for each sample (bars on x-axis), sectioned by geographic region. The three K populations identified aligned perfectly with the clustering in the NJ tree; K=1 with *Mf*, K=2 with *Mn* and K=3 with Peninsular as per the colour-coding

Despite substantial genetic divergence between the *Mf* and *Mn* clusters (mean F_ST_ = 0.2), there is also substantial geographic overlap (*Figure 2.B*) and evidence of shared ancestry (>1% ancestry to two or more groups) amongst 6.6% (n=10/152) of infections (*Figure 2.C*). This observation extends beyond the newly sequenced Sabah samples and to those previously reported in neighbouring Sarawak, with the separate genomic *Mf* and *Mn* clusters and several samples of *Mf* and *Mn* ancestry being identified in both geographic locations. *P. knowlesi* infections with shared ancestry originated from the Sarikei and Betong districts in the state of Sarawak and four of the newly sequenced Sabah infections (from Papar, Ranau and Kota Marudu districts). Although several Malaysian-Borneo *P. knowlesi* infections had evidence shared ancestry, samples from Peninsular-Malaysia and the district of Kapit in Sarawak (both *Mn* and *Mf*) are descendants of single ancestral populations (*Figure 2.C*).

### Greater genetic diversity within the *Mf* than *Mn* and *Peninsular* clusters

Since malaria parasites are recombining organisms, neighbour-joining analysis can miss recent connectivity between infections where outcrossing has taken place. To further elucidate the relatedness between isolates, both within and across clusters, we performed identity-by-descent (IBD) analysis on the 152 low complexity infections. In IBD analysis, genomic segments are characterised as identical by descent in pairwise comparisons when identical nucleotide sequences have been inherited from a common ancestor. At IBD thresholds under ∼30%, three relatively distinct, large infection networks were observed, largely consistent with the *Mf*, *Mn* and *Peninsular* groupings defined by IBS analyses (*Figure 3*). The *Mf* and *Mn* clusters shared a small degree of connectivity (∼3%), with *Mf* isolates from Sabah (PK_SB_DNA_028 – Papar and PK_SB_DNA_053 – Kudat) connecting to a selection of *Mn* isolates, also from Sabah. The *Mf* cluster had the highest genetic diversity, with a median IBD of 7.0%, and as such, we see the associated network break down at a relatively low threshold of 11.8%. In contrast, the *Mn* (median IBD = 0.5) and *Peninsular* (median IBD = 0.3) clusters maintain tight networks at 30.4% IBD, reflective of more recent common ancestry and a greater number of shared haplotypes, and in turn, lower transmission intensity. To evaluate whether the high IBD values in *Mn* and *Peninsular* clusters were not inflated by the SNPs used (i.e. being biased by strong population structure), we trialled multiple filtering combinations and re-calculated the median IBD values for comparison, confirming our initial findings (Supplementary Table 3).

**Figure 3.**
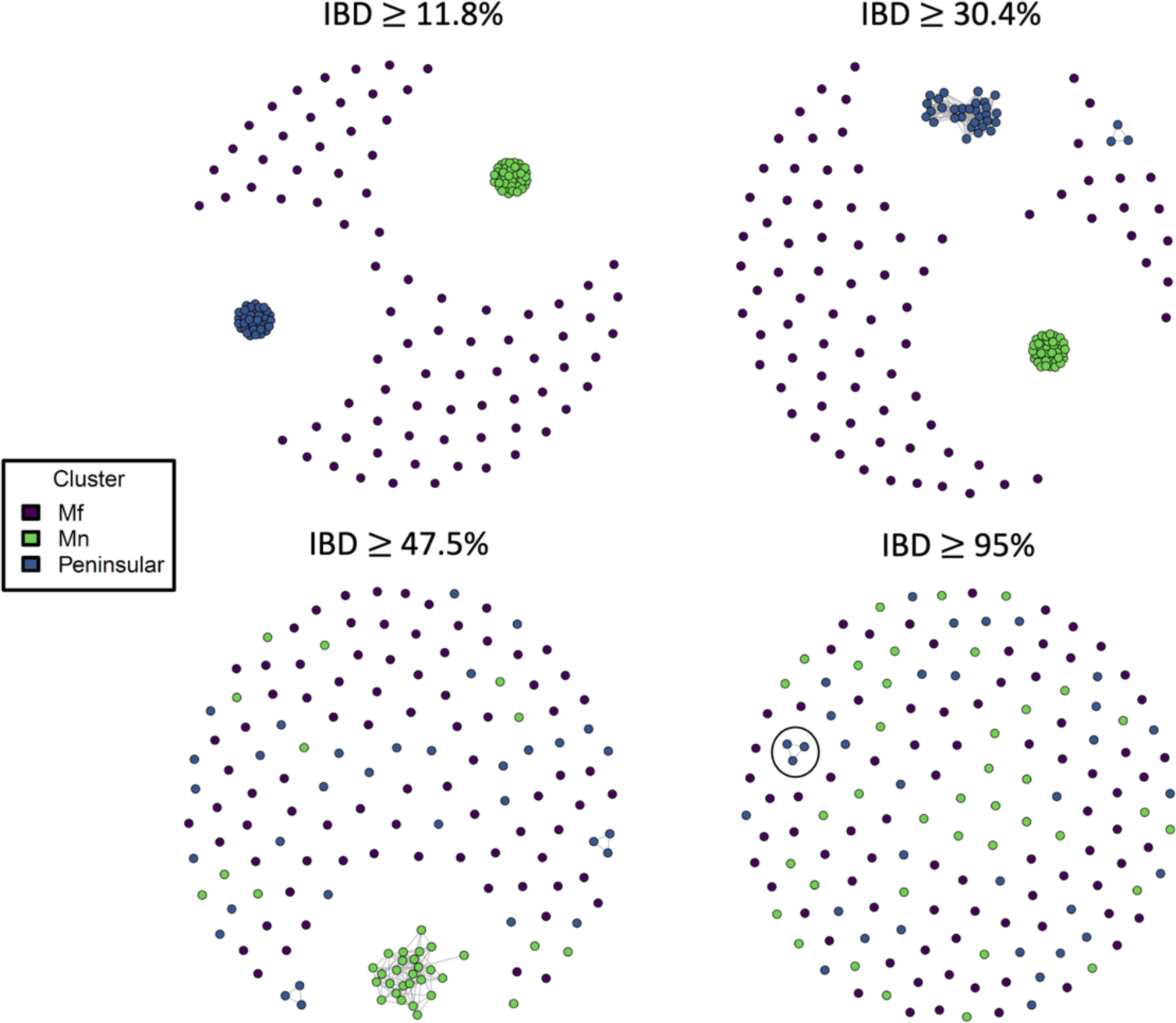
Identity by decent (IBD)-based analysis reveals greater relatedness amongst *Mn* and *Peninsular* than *Mf* samples. Each circle reflects an infection, colour-coded by genomic clustering group, and line lengths reflect relatedness (shorter lines reflect greater relatedness) at the given connectivity threshold of minimum IBD. Where two circles are not connected by a line, the estimated IBD between those infections was below the given threshold. The distance between infections (circles) that are not connected by lines does not reflect the relatedness between those infections. IBD thresholds were determined by halving 95% and every subsequent threshold, with additional values included to show the spectrum of the breakdown in relatedness across the population clusters. The three samples from Peninsular Malaysia with >95% IBD represent laboratory-based strains from the 1960s that have been passaged through macaques (SRR2222335, SRR2225467 & SRR3135172).

### Substantial evidence for introgression between *Mn* and *Mf* clusters

Previous studies have described the occurrence of chromosomal-segment exchanges between the *Mn* and *Mf* subpopulations, suggesting that they are not genetically isolated (15). We therefore sought evidence for introgression events in our large collective cohort, and specifically, in the previously underrepresented Sabah. Comparisons of genetic distance between 10kb sliding windows in individual *P. knowlesi*-infected samples and different clusters reveal evidence of substantial genetic exchanges across genomic clusters, chromosomes, and geographical regions (Figure 4 & Supplementary Table 4). The degree of introgression, represented by the number of introgressed windows identified in a sample, also varied between all of the above-mentioned features.

**Figure 4.**
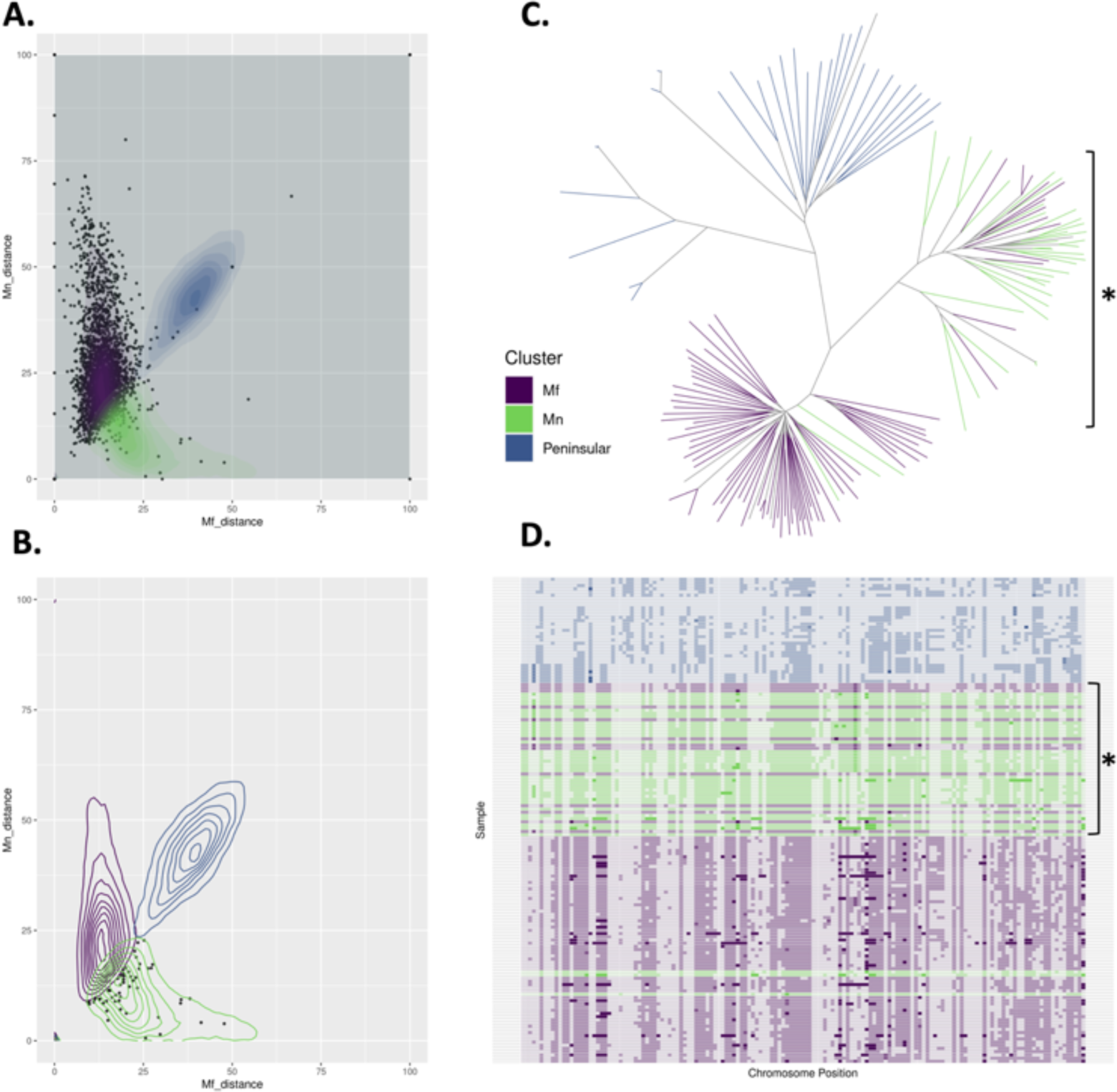
Determination of introgressed windows between the *Mf* and *Mn P. knowlesi* clusters. A: Dot and contours plot describing potential introgression events within an *Mf* sample, where x and y axes represent the genetic distance of the sample to the *Mf* and *Mn* clusters, respectively. Genetic distance is the proportion of mismatched SNPs per sliding window (10kb) when comparing the called allele in the sample to the major allele for a cluster at each position. The contours represent the density of genetic distances for the three clusters. B: Dot and contours plot of the same sample above, subset to those windows deemed to be introgressed from the *Mn* cluster. Possible introgression events are sliding windows that fall outside the major contours of the samples own cluster and within the major contours of another, representing greater similarity in genetic distance to the other cluster. C: Unrooted neighbour-joining tree based on identity by state (IBS) of *window 1504* on chromosome 08 (950000-959999) and overlapping the PKA1H_080026000 gene (encodes the oocyst capsule protein). The *Mf* samples/branches clustering within the *Mn* branches (depicted by asterisk) provides further evidence that introgression of this window has occurred in these samples. D: SNP barcode plot of *window 1504* on chromosome 08 (950000-959999) showing greater genetic similarity between several *Mf* samples (depicted by asterisk) and the *Mn* cluster, where the colours are reflect those in the legend above, and the alpha represents the allele.

Of the 152 individual *P. knowlesi* samples analysed, 71.1% (111/152) had two or more windows considered to be introgression. Approximately 29.5% (n=46/152) of samples demonstrated a high level (>5 introgressed windows) of introgression. Within the subset of newly generated genomes from Sabah, 82.7% (n=43/52) samples had two or more windows of introgression, including 20 at high levels. The *Mf* cluster had a higher median number of introgressed windows per sample (median 5, IQR ±9.27) compared to *Mn* (median 1, IQR ±0.88). Candidate windows with introgression events appeared across ten chromosomes for *Mf* and six chromosomes for *Mn*. For the *Mf* cluster, chromosomes 8 (n = 35) and 11 (n = 21) had the greatest number of candidate windows, and for *Mn*, all six chromosomes contained a single window. The district of Betong in Sarawak had the highest median number of introgressed windows per individual *P. knowlesi* sample (median 29, IQR ± 12.2) followed by the district of Papar in Sabah (median 19, IQR ± 25.46). 85% percent of Betong samples (n =12/14) and 50% of Papar samples (n=1/2) had high levels of introgression, with all but one Betong sample from the *Mf* cluster. The *Mf* isolate from Papar with high levels of introgression (*PK_SB_DNA_028*), had the greatest number of candidate windows overall (n = 37), followed by ten *Mf* samples from Betong that had greater than 20 introgressed windows (Supplementary Table 4).

Several candidate introgressed regions identified in the *Mf* clusters overlap genes involved in host interactions. Within Sabah, the most common candidate window (window 1504, chromosome 11: 2080000 - 2089999), observed in both Sabah (n = 17) and Sarawak (n = 7) isolates, overlaps a gene encoding for the parasite DNA repair protein RAD50 (PKA1H_110050600), which may aid survival within the host (26). Focusing on the top 10 most abundant windows in both Sabah and Sarawak, several other genes encoding for proteins essential for survival or invasion in the human, macaque or mosquito hosts were also identified to overlap candidate windows (Table 1). Amongst windows more prevalent in samples from Sarawak were several genes that encode for mosquito-related proteins. This includes the oocyst capsule protein (PKA1H_080026000), CPW-WPC family protein (PKA1H_080026200) and the microneme-associated antigen (PKA1H_080031400). The inclusion of the new Sabah isolates expands the distribution of the introgression event associated with the oocyst-expressed *cap380* gene, previously only observed in Betong, Sarawak (15, 18). The oocyst capsule protein is essential for the maturation of ookinete into oocyst in *P. berghei* and is assumed to assist in immune evasion in mosquito hosts (27). The CPW-WPC proteins are zygote/ookinete stage-specific surface proteins and appear to be involved in mosquito-stage parasite development (28) and micronemes are critical for host-erythrocyte invasion (29). The number of candidate regions in *Mn* was minimal (n = 6), with even fewer involved in host interactions. The exception being window 807 on chromosome 08 (760000–769999), overlapping the PKA1H_080021900 gene, which is essential for erythrocyte invasion in *P. falciparum* (30).

**Table 1.** Genes overlapping the ten most common candidate regions for introgression in *Mf* and *Mn*. Description of gene name, position, encoded protein, and the isolate counts and proportions for the two Malaysian Borneo states.

| Window | Gene | Chromosome: start - end | Protein | Sabah (n [%]) | Sarawak (n [%]) |
| --- | --- | --- | --- | --- | --- |
| 1504 | PKA1H_110050600 | 11: 2080000 – 2089999 | DNA repair protein RAD50 | 17<br>[27.78] | 7<br>[10.78] |
| 1216 | PKA1H_100020600 | 10: 723,246 – 723,968 | Apicortin | 8<br>[14.81] | 1<br>[1.54] |
| 1216 | PKA1H_100020800 | 10: 720000 – 729999 | F-actin-capping protein subunit beta | 8<br>[14.81] | 1<br>[1.54] |
| 1223 | PKA1H_100022400 | 10: 790000 – 799999 | NLI interacting factor-like phosphatase | 8<br>[14.81] | 1<br>[1.54] |
| 1236 | PKA1H_100025500 | 10: 923,327 – 924,091 | Orotate phosphoribosyltransferase | 8<br>[14.81] | 2<br>[3.08] |
| 1359 | PKA1H_110018600 | 11: 629,218 – 635,353 | Patatin-like phospholipase | 8<br>[14.81] | 7<br>[10.77] |
| 1309 | PKA1H_110008500 | 11: 138,605 – 141,595 | Secreted ookinete protein | 6<br>[11.11] | 9<br>[13.85] |
| 1309 | PKA1H_110008100 | 11: 128,971 – 130,944 | ABC transporter E family member 1 | 6<br>[11.11] | 9<br>[13.85] |
| 31 | PKA1H_010010000 | 01: 299,231 – 301,798 | DNA mismatch repair protein<br>MSH2 | 6<br>[11.11] | 3<br>[4.62] |
| 826 | PKA1H_080026000 | 08: 940,469 – 950,653 | Oocyst capsule protein | 1<br>[1.85] | 10<br>[15.38] |
| 826 | PKA1H_080026200 | 08: 954,493 – 956,175 | CPW-WPC family protein | 1<br>[1.85] | 10<br>[15.38] |
| 826 | PKA1H_080026300 | 08: 956,849 – 958,271 | Protein transport protein SEC22 | 1<br>[1.85] | 10<br>[15.38] |
| 827 | PKA1H_080026700 | 08: 968,445 – 976,496 | ABC transporter I family member<br>1 | 1<br>[1.85] | 10<br>[15.38] |
| 854 | PKA1H_080031400 | 08: 1,229,556 –<br>1,230,530 | Microneme associated antigen | 3<br>[5.56] | 10<br>[15.38] |
| 819 | PKA1H_080024200 | 08: 878,480 – 880,896 | ATP-dependent RNA helicase<br>DDX6 | 1<br>[1.85] | 9<br>[13.85] |

**Table 2.** Summary of identified drug resistance orthologue mutations. . Drug resistance orthologues from *P. vivax* and *P. falciparum* (none identified from *P. falciparum*) identified in this *P. knowlesi* dataset. ^+^Designates the number of amino acid positions the identified *P. knowlesi* mutation is from the corresponding *P. vivax* orthologue mutation position (proximal mutations may retain the potential to cause similar downstream effects). AA: amino acid change; Pk: *P. knowlesi*. Pv: *P. vivax*.

| Gene | *Pv* gene ID | *Pk* gene ID | AA – *Pv* | AA – *Pk* | Anti-malarial |
| --- | --- | --- | --- | --- | --- |
| *pvdhps* | PVP01_1429500 | PKA1H_140035100 | C422W | G422S | Sulphadoxine |
| *pvdhfr* | PVP01_0526600 | PKA1H_050015200 | N273K | N272K<sup>+++</sup> | Pyrimethamine |
| *pvdhfr* | PVP01_0526600 | PKA1H_050015200 | N273K | N272S<sup>+++</sup> | Pyrimethamine |
| *pvmrp1* | PVP01_0203000 | PKA1H_140054000 | I1620T | C1908S<sup>+</sup> | Multi-drug |
|  |  |  | C1018Y | V1283I<sup>+</sup> |  |
|  |  |  | K542E | N620K<sup>++</sup> |  |
|  |  |  | R259T | G316D<sup>+++</sup> |  |
|  |  |  | R259T | G316S<sup>+++</sup> |  |
|  |  |  | T234M | M251K<sup>++</sup> |  |
|  |  |  | T234M | E250G<sup>+++</sup> |  |

### Ecological pressures driving introgression

In order to evaluate whether the cluster distribution or the introgression events are associated with ecological changes that might impact either macaque host or vector adaptations, we collated satellite-based surrounding forest fragmentation data and mosquito vector habitat suitability for 37 *P. knowlesi* samples in Sabah where village locations could be obtained (Supplementary Figures 2 & 3). These samples included 29 (78.4%) with two or more windows where introgression was observed, and 13 (35.1%) with high introgression (>5 windows).

Firstly, we performed univariate regression analyses of the *P. knowlesi* genomic clusters against proportional forest cover, intact forest perimeter-area ratio and *Anopheles* Leucosphyrus Complex mosquito vector habitat suitability metrics, with no statistically significant associations. Secondly, univariate regression analyses (optimal model as determined by AIC comparisons) suggested a limited relationship between two candidate windows for introgression, 859 (chromosome 08: 1280000 – 1289999) and 1236 (chromosome 10: 920000 – 929999) and the intact forest perimeter-area ratio and mosquito vector habitat suitability, respectively (Supplementary Tables 5 & 6). The introgression of window 859, which contains no identifiable genes on *PlasmoDB* and was identified in three Sabah and ten Sarawak isolates, was positively associated with intact forest perimeter-area ratio (*ꭓ^2^* = 6, df = 1, p = 0.02, r^2^ = 0.69). The introgression of window 1236, which was identified in eight Sabah and two Sarawak isolates, was negatively associated with the predicted mosquito vector habitat suitability (*ꭓ^2^* = 8.17, df = 1, p < 0.01, r^2^ = 0.71). Genes overlapping this region include two encoding for unknown proteins, one encoding for ras-related protein Rab-1B and another for orotate phosphoribosyltransferase (*Table 1*).

### Greater relatedness within that between state-level subpopulations

It was hypothesised that *P. knowlesi* clinical infections derived from the separate states of Sabah and Sarawak in Malaysian Borneo are likely to have distinct genetic ancestry due to factors such as differences in the primary *Anopheles* Leucosphyrus Group mosquito vector species and other large-scale environmental features that may have restricted historical gene flow (31). To test this, we leveraged the newly sequenced genomes to perform additional analyses on a subset of the data comprising isolates from Sabah and Sarawak. We examined the potential impact of geographical regions on population structure and genetic relatedness within each of the separate *Mf* and *Mn* clusters (samples subset to clusters for analysis). IBD analyses of *Mf* and *Mn* subsets suggest that most samples have greater connectivity within their respective states (*Figure 5*).

**Figure 5.**
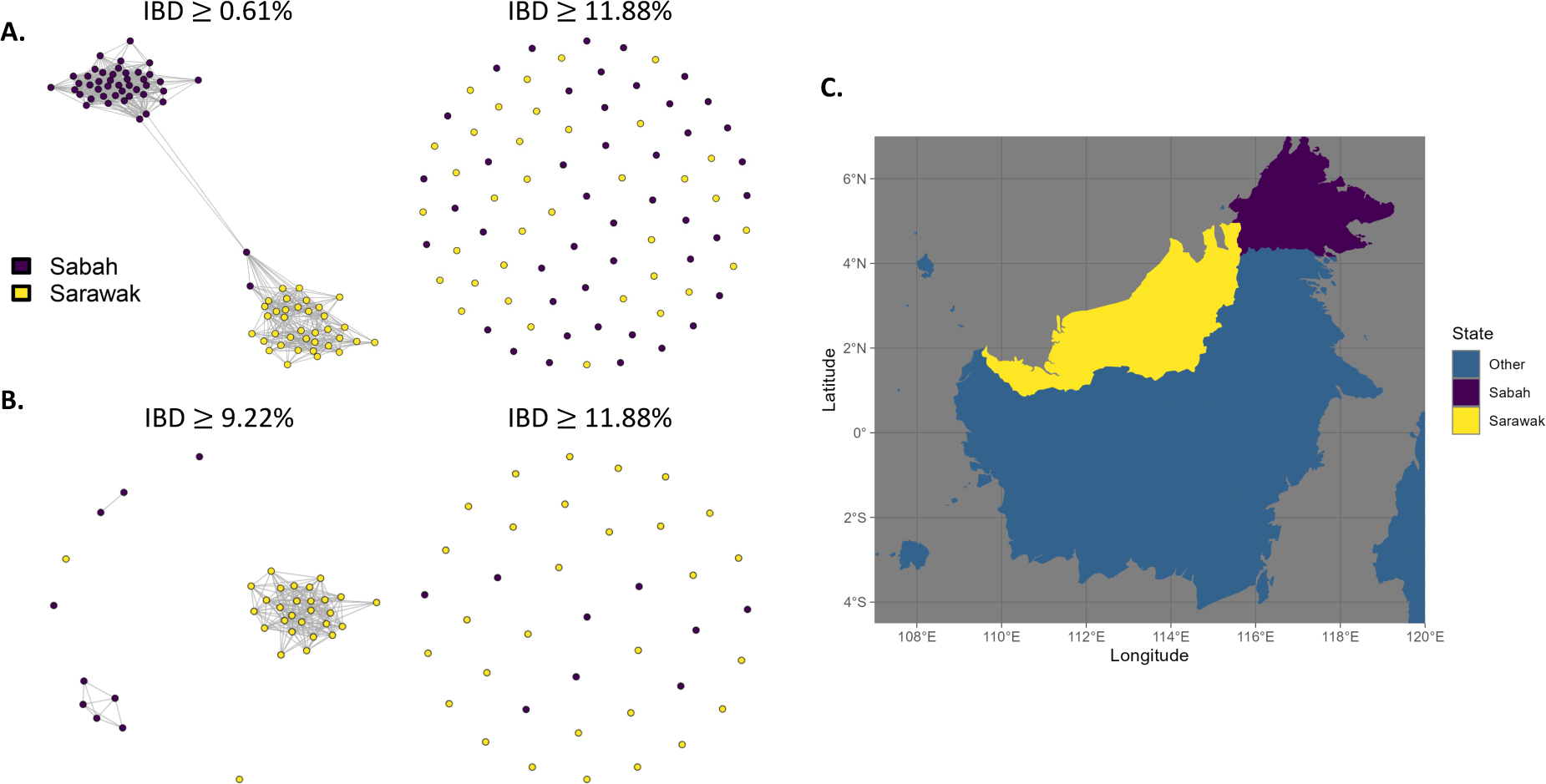
Infection connectivity is partly driven by geography at the state-level administrative boundary. A & B: Identity by decent (IBD)-based cluster network illustrating the distant relatedness for samples within *Mf* (A) and *Mn* (B) clusters collected in two adjacent states; Sabah and Sarawak, at different cut-offs for the proportion of IBD in a paired comparison. C: Map of East Malaysia on the island of Borneo with colours representing the two states being compared in the IBD analysis.

Two Sabah samples within the *Mf* cluster had a high degree of connectivity with Sarawak samples (*Figure 5A*). The samples (PK_SB_DNA_028 and PK_SB_DNA_053) are *P. knowlesi* infections collected from residents of the Papar and Kudat districts in Sabah and were likely not collected from a proximal district to Sarawak, although no travel history was obtained. Despite this, both samples exhibit substantial patterns of introgression, in a similar pattern (the same or proximal windows) to that seen in the samples from Sarawak with which they share a relatively high degree of IBD connectivity (Supplementary Table 7). Thus, recent genetic transfer events arising independently across multiple districts and states may explain the high degree of IBD between these samples.

### Population differentiation within *Mf* and *Mn* clusters of geographic subpopulations

Given that sampling sites of the *P. knowlesi* geographic subpopulations are isolated by several hundred kilometres, with spatially heterogenous environmental pressures, we performed genome-wide scans for differentiation between the geographic subpopulations within *Mf* and *Mn* subsets. Genome-wide scans within *Mf* highlighted several regions of significant differentiation across the genome, appearing as peaks of multiple tightly clustered windows of high F_ST_ against a background of low differentiation (mean F_ST_ = 0.007, *Figure 6A*). The most notable peaks within the *Mf* cluster were observed on chromosomes 8, 11 and 12, with the peak on chromosome 8 spanning a cluster of introgressed windows identified in our introgression analysis. This includes the introgressed window containing the gene encoding for the oocyte capsule protein. This window appeared to be introgressed across several Sarawak isolates and one of the Sabah isolates (PK_SB_DNA_028) shown to maintain connectivity to Sarawak in the cluster specific IBD analysis (*Figure 5A*). A complete list of genes found in the peak can be found in the Supplementary Tables 8 and 9. Unfortunately, due to substantial noise, it was not possible to appropriately identify peaks for the *Mn* cluster (mean F_ST_ = 0.036, *Figure 6B*).

**Figure 6.**
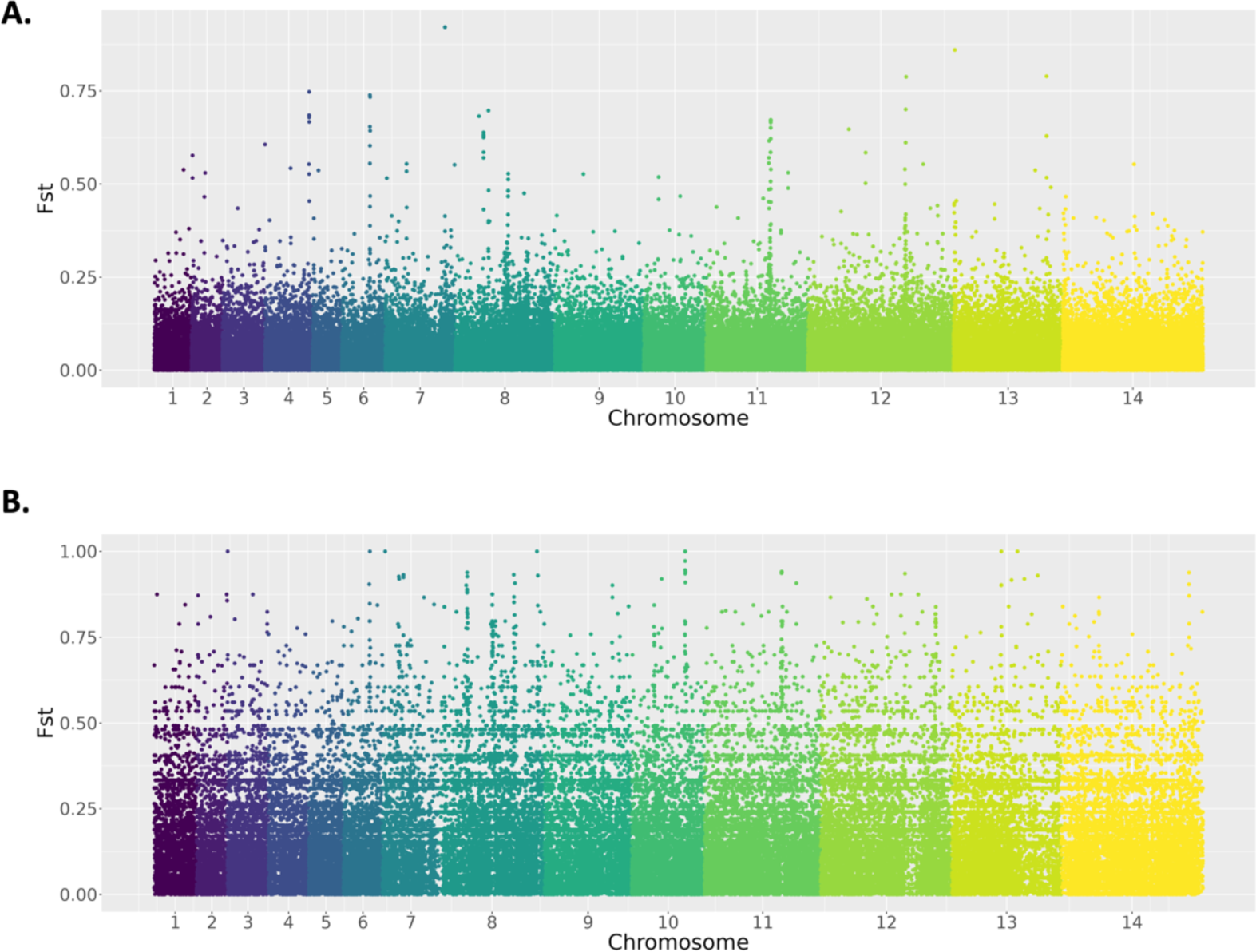
Genetic differentiation reveals candidate adaptations. Genome-wide scans of differentiation between Sabah and Sarawak subpopulations within the (A) *Mf* and (B) *Mn* clusters using the between-population fixation index (F_ST_). Only the *Mf* cluster shows clear differences in diversity with peaks of differentiation clear at several chromosomes (most notably on chromosomes 8, 11 and 12), whilst the *Mn* cluster has substantial ‘noise’ across the genome, with high levels of differentiation across the genome.

### Investigation of antimalarial drug resistance candidates in *P. knowlesi* orthologues

The presence of antimalarial drug resistance determinants in *P. knowlesi* infections could be considered a surrogate marker of human-human transmission given the absence of drug pressure in the macaque hosts and fitness costs that are often associated with resistance-conferring alleles (32). We therefore investigated the prevalence of non-synonymous variants in *P. knowlesi* orthologues of genes that have previously been associated with *P. falciparum* and *P. vivax* resistance to antimalarial drugs (33, 34). Within low complexity infections (n=152), six non-synonymous variants were detected within the *P. knowlesi* orthologue of *pvdhps* (PKA1H_140035100),), which may be linked to sulphadoxine resistance (34). The most common variants occurred at codon Y308H (58.4%) and K66E (11.9%). A G422S variant was present in 4.3% of samples overall, although was found exclusively in 25% of isolates from Peninsular Malaysia. Similar to previous work (32), 13 non-synonymous mutations were also detected within the *P. knowlesi* orthologue to *pvdhfr* (PKA1H_050015200). This includes 17 samples with greater than one mutation, and two samples with three mutations. Dihydrofolate-reductase mutations, associated with resistance to pyrimethamine, arise readily in both *P. falciparum* (35) and *P. vivax* (36, 37). The most common mutations were at codon N272S (97.9%) and E262D (22.7%), with N272S occurring 3 amino acid positions from the *P. vivax* orthologue. Six mutations were also observed exclusively in isolates from Peninsular Malaysia, with a mean frequency of 10.6%. Lastly, several non-synonymous mutations were also observed in the PKA1H_100015100 gene, resulting in several amino acid changes near those observed in *P. vivax*, and associated with the multidrug resistance protein 1 (*pvmrp1*). This includes seven amino acid substitutions that occur <3 amino acids from non-synonymous mutations present in *pvmrp1* potentially linked to primaquine failure in *P. vivax* liver-stage infection relapse. Despite the proximity of these non-synonymous mutations to known drug-resistance orthologues, homology modelling would be required to explore whether they would be translated into drug-resistance phenotypes.

## Discussion

In this work, we expand the analytical approach used to understand the previously described population genetics of *P. knowlesi*, including the use of an additional 52 new whole genomes from the previously under-represented state of Sabah, a major focus of *P. knowlesi* transmission in Malaysia, and report several new findings. First, using plots of within-isolate non-reference allele frequencies, we identify and describe a high prevalence of both recombinant co-transmission and additive super-infections of multiple *P. knowlesi* lineages in humans. Second, using an IBD-based connectivity analysis, we have identified the emergence of distinct geographical subpopulations within the *Mf-* and *Mn*-associated clusters. Furthermore, we characterise evidence of widespread introgression events between the *Mf-* and *Mn*-associated clusters in Sabah, which may be linked to differentiation of the geographical subpopulations. Building on this, we conducted an exploratory approach to integrating ecological and genomic data, using *P. knowlesi* sample locations and surrounding forest fragmentation and mosquito vector habitat suitability data. Evaluating genomic cluster distribution or introgression events within this framework provided preliminary insights into several possible associations with ecological features that may impact either host or vector adaptations. Lastly, we further characterised the presence of non-synonymous mutations across *P. knowlesi dhps, dhfr* and *mrp1* gene orthologues associated with antimalarial drug resistance in other *Plasmodium* species, despite the primary zoonotic transmission mode of *P. knowlesi* likely limiting historical drug selection pressure and the development of drug resistant clinical phenotypes (38, 39).

Within-host *Plasmodium* diversity is often used as a proxy for understanding transmission intensity in endemic areas, with superinfections expected to be more common in high-transmission settings where there is typically a large diversity of parasites that can be transmitted to humans through multiple inoculated mosquito bites (40, 41). Polyclonal infection can also arise through co-transmission, where a single mosquito inoculum comprises multiple (generally related) parasite strains (11). The prevalence of apparent superinfections observed in Sabah is lower than frequencies observed in *P. faciparum* or *P. vivax* infections in moderate to high endemic settings (42, 43). However, it needs to be considered that the proportion of *P. knowlesi* superinfections described here is only reflective of the human host population and may be higher in the natural macaque hosts (44). Infection complexity in *P. knowlesi* is complicated by the zoonotic nature of *P. knowlesi* transmission, with multiple underlying parasite, host and epidemiological factors potentially influencing the establishment of successful erythrocytic replication of multiple inoculated *P. knowlesi* strains within humans causing symptomatic disease. This includes the presence of appropriate parasite human red blood cell invasion mechanisms (45), the degree of transmission intensity within the adjacent macaque host reservoir, and underlying environmental drivers impacting mosquito bionomics and host biting preferences.

Inter-infection diversity can also be exacerbated by recombination between genetically distinct parasites within the mosquito host (11). The result of such recombination is the highly related clones observed in the NRAF plots, which most probably derive from the co-transmission of polyclonal infections originating in the macaque hosts. Being a natural reservoir for multiple zoonotic *Plasmodium* species, macaques have been shown to be co-infected with up to five simian *Plasmodium* species (44) and multiple *P. knowlesi* clones (46, 47). Alternatively, the isolates in which we see distinct clones are more likely to be examples of superinfections, as multiple clones originating from the same macaque host would have undergone recombination during reproduction in the mosquito host. Although widespread naturally-occurring human-to-human transmission is not evident for *P. knowlesi* to date (48), and the risk of *P. knowlesi* infection remains relatively low for most populations, high at-risk occupational groups have been identified, such as forestry and plantation workers (9, 10). These jobs place individuals in closer proximity to the vector and reservoir hosts for longer durations, including at peak mosquito biting times, putting them at a greater risk of infection (49), and potentially, superinfections. It is possible that the five individuals who were unfortunate enough to be inoculated multiple times belong to a high-risk demographic. Future work should look to integrate larger epidemiological and targeted genomic datasets to appropriately answer such questions.

Consistent with previous work, both neighbour-joining and IBD-based cluster analyses reveal three predominant *P. knowlesi* genomic clusters in Malaysia. Of the newly explored *P. knowlesi* isolates from Sabah the overwhelming majority were within the established *Mf* cluster, with only nine isolates from the *Mn* cluster. The large number of *Mf* cluster infections are consistent with reports of a higher prevalence of *Mf-*derived human malaria cases across Malaysia (22), as well as the more restrictive ecosystem distribution of *M. nemestrina*, which prefer intact forest areas (50). Evidence of higher transmission intensity and more diverse populations of *Mf* infections are evident in the low IBD median for the cluster, with low population IBD characteristic of highly endemic *Plasmodium* species populations due to a lower level of inbreeding. In contrast, the higher median IBD values for the *Mn* cluster suggests a greater degree of parasite relatedness and inbreeding, with many of the pairwise percentage-IBD values indicative of siblings or half siblings. However, as the median IBD reduces substantially when down sampling to only the *Mn* cluster, we suggest the inflated values may be a product of the strong population structure (genetic distance between clusters) of *P. knowlesi* (14), as the algorithm used is designed for analysis in a single population. The comparatively larger amount of genetic differentiation within *Mf* isolates may have resulted from the broader range of ecosystems that the *M. fascicularis* inhabit, the larger troop sizes, and a greater ability to adapt to changing ecosystems associated with land clearing and agricultural activities. This includes cropland, wetland, vegetation mosaics and peri-urban areas (50). The high genetic diversity seen in the *Mf* cluster may also impact future malaria control efforts, as this feature has the potential to increase the parasites’ ability to adapt in response to ongoing environmental selection pressures, leading to increased zoonotic transmission through improved host invasion and survival.

Deforestation and agricultural expansion have significantly impacted macaque and *Anopheles* habitats and behaviour in *P. knowlesi* endemic areas (49, 51, 52), which are hypothesised to influence recent parasite genetic exchange events seen in human clinical infections (14). In this work, regression analyses identified significant associations between two introgressed regions of the genome and both forest fragmentation (perimeter-area ratio) and predicted habitat suitability of the *Anopheles* Leucosphyrus Complex-mosquito vector. Supporting the previous stated hypothesis. Several of the introgressed windows contain genes encoding for proteins critical to invasion and survival in the human and mosquito hosts. Examples of *P. knowlesi* orthologous proteins identified as critical to parasite replication in mosquitos in *P. falciparum* are the microneme-associated antigen, which binds to the erythrocyte surface receptors to facilitate erythrocyte entry (53), and the previously identified oocyst-expressed *cap380* gene, essential for transmission of the parasite during the vector stage (27), and previously only identified in Betong, Sarawak (14). Relevant to human infection, one example is the Apicortin protein, which maintains the stability of the parasite cytoskeletal assemblies in *P. falciparum* and *P. vivax*, and in turn, is important for replication, motility, merozoite invasion of host erythrocytes and survival (54, 55). Several other vector and human-related genes were identified to be present in recent introgression events, suggesting strong evolutionary selective pressure on the parasites driven by both hosts. Furthermore, the occurrence of these events across large geographic distances (assuming a lack of self-reported travel history) suggests introgression events could be arising independently through similar environmental drivers, such as deforestation and changing vector populations. However, this integrated genomic and spatial analysis is limited in part by the lack of exact temporal alignment between *P. knowlesi* infection collection timepoints and the environmental data. Additionally, as these analyses were performed on a relatively small subset of isolates, with a small number of previously validated landscape metrics selected at a single spatial scale (56), future work should include a larger sample set and broader systematic approach to selection of landscape classification variables encompassing temporal land use change.

The spatial distribution of *P. knowlesi* infection sampling sites across Malaysian Borneo occurred in two distinct large-scale geographical areas. Although the macaque hosts are reported to be the primary driver of *P. knowlesi* genetic population structure, there are likely to be other underlying environmental determinants that vary across spatially heterogenous landscapes. To explore the impact of geographical differences in environment (without constraints imposed by defined population structure), we evaluated *P. knowlesi* infections separately within the *Mf* and *Mn* clusters by sub-setting the isolates to their respective clusters and replicating analyses within these subsets, making comparisons between Sabah and Sarawak subpopulations. As expected, within both the *Mf* and *Mn* clusters there is a higher degree of relatedness between samples collected in closer proximity, albeit less so when compared to the three major genomic clusters. The exception being two samples from Sabah with high levels of introgression that clustered closer to those from Sarawak. The high degree of IBD between these samples may be explained by recent genetic transfer events arising independently across multiple districts and states. However, given the lack of corroborative travel history for the infected individuals, we are unable to exclude the possibility that these infections were also obtained in Sarawak.

The stronger relatedness between geographically proximal samples suggests that in addition to the macaque hosts, other ecological pressures involved in zoonotic *P. knowlesi* transmission cycles are impacting the genomes. This was confirmed through F_ST_ analysis, specifically of the *Mf* cluster. Although many of the genes spanning outlier regions in *Mf* were unknown proteins, many of these same genes were also identified through the introgression analysis, such as the oocyst-expressed *cap380* gene on chromosome 08. Thus, the intermediate mosquito host may be a driver of subpopulation-level differences, especially considering that the species of the primary *Anopheles* Leucosphyrus Complex group mosquito vector differs across regions (57). Human land use change has also led to adaptive behaviors, changes in larval breeding sites, and host biting preferences in the primary vector in Sabah, *A. balabacensis* (58). To better resolve these signals of selection, particularly for the *Mn* cluster, future work may be aided by using the cluster-specific reference genomes published by Oresegun *et al.* 2022 (59).

Despite significant progress towards the elimination of the major causes of human malaria, *P. falciparum* and *P. vivax*, antimalarial drug resistance remains a major public health threat (60). However, as *P. knowlesi* transmission appears to remain exclusively zoonotic (2), drug-resistance mutations are less likely to emerge. Despite this, we observed several SNPs in orthologue genes previously highlighted as potential drug resistance markers, including dihydropteroate synthase (*dhps*), dihydrofolate reductase (*dhfr*) and the multidrug resistance-associated protein (*mrp1*). *dhfr* and *dhps* may be associated with pyrimethamine and sulphadoxine resistance, respectively, which were previously used in combination for the treatment of *P. falciparum* malaria in Malaysia from the 1970s, and up to recent years for household contacts of malaria outbreaks due to any *Plasmodium* species (61). The variant causing the highly prevalent N272S mutation within the *dhfr* gene, is only present in isolates from lab-adapted lines. The A allele is observed in four reference isolates that have been passaged through macaques, as well as the A1H1 reference strain (62). However, the remaining isolates, along with the PKNH reference genome (63), all contain the G allele. It appears that the G allele has displaced A and is now fixed in the population. For the other mutations, it is possible that sulphadoxine-pyrimethamine inadvertently and successfully treated a proportion of historical *P. knowlesi* infections misidentified by microscopy (64).

The current recommended blood-stage treatment for all *Plasmodium* species in Malaysia causing uncomplicated malaria, including *P. knowlesi*, is with artemisinin-combination therapy (65, 66), meaning no ongoing drug selection pressure for sulphadoxine-pyrimethamine remains present (67). We have previously shown that *dhfr* mutations in *P. knowlesi* are unlikely to mediate drug resistance and are therefore unlikely to have been selected by past pyrimethamine exposure (32). Furthermore, others have previously failed to identify mutations associated with *dhps* in *P. knowlesi* (68). Given the lack of both widespread human-to-human transmission and the use of sulphadoxine-pyrimethamine in *P. knowlesi* treatment regimes, the cause and functional implications of these observations are unclear. It is likely that these mutations are due to the polymorphic nature of these genes rather than drug selection pressure. In addition, although variants linked to *mrp1* have previously been observed (16), with unknown consequences, it is also unclear if the SNPs observed in this work would have structural implications in the resulting protein.

The addition of 52 high-quality whole genomes from a major regional focus of transmission in Sabah, Malaysia broadens our understanding of the complex and evolving genomic landscape of *P. knowlesi*. We identify both *P. knowlesi* superinfections and co-transmitted infections. Through IBD-based connectivity analysis, we demonstrate the emergence of regional geographical subpopulations, which may be linked to recent introgression events between the *Mf-* and *Mn*-associated clusters in Sabah. Furthermore, we show that introgression events may be associated with ecological changes reflective of either host or vector adaptations, and lastly, we identify several non-synonymous mutations across the *dhps, dhfr* and *mrp1* drug-resistant genes. Changing ecosystems in Malaysia, driven by anthropogenic deforestation and agricultural activities are bringing humans into closer proximity to the reservoir and vector hosts. Our data suggests that these underlying drivers of *P. knowlesi* transmission are also consistent with the genetic changes seen in *P. knowlesi* populations. Taking lessons from *P. falciparum* and *P. vivax*, continuing to grow our understanding of *P. knowlesi* genetics with large scale genetic surveillance will be invaluable in guiding future public health surveillance and control strategies.

## Methods

### Sample collection and preparation

We used a combination of newly generated *P. knowlesi* whole genome sequencing data (n = 94) (69) and archived FASTQ files from samples (n = 108) from *P. knowlesi*-infected patients in Malaysia. Newly processed samples were collected as part of prospective clinical studies conducted through the Infectious Diseases Society Kota Kinabalu Sabah-Menzies School of Health Research collaboration from 2011 to 2016 across multiple hospital sites in Sabah (4, 5). Patients of all ages presenting with microscopy-diagnosed malaria were enrolled following informed consent. *P. knowlesi* infections were confirmed through validated PCR (70, 71, 72) and parasitemia quantified by expert research microscopists. These 94 clinical isolates underwent Illumina whole genome, paired-end sequencing (150bp), with library preparation conducted using the NEBNext® Ultra™ IIDNA Library Prep Kit (from New England BioLabs Inc., Cat No. E7645). Ethical approval was obtained from the medical research ethics committees of the Ministry of Health, Malaysia and Menzies School of Health Research, Australia. A further 108 samples from a broader geographic range within Malaysia were downloaded from the National Center for Biotechnology Information (PRJEB33025, PRJEB23813, PRJEB1405, PRJEB10288 & PRJN294104) (14, 20) (Supplementary Table 1).

### Read mapping, variant discovery and genotyping

Variants were detected using a modified version of a previously described workflow (73). Raw reads were processed using FastQC and cutadapt (74) to determine quality, with subsequent filtering and trimming of reads. The Burrows-Wheeler Aligner (bwa) was then used to map reads to the PKA1-H.1 reference genome (23). BAM pre-processing steps were applied using Picard version 2.26.1 and the Genome Analysis Toolkit (GATK) version 3.8-1-0 (75). Notably, two steps in the GATK best-practices pipeline (base recalibration and indel realignment) require a set of high-quality known variants. As no such variant lists existed for *P. knowlesi*, we generated a conservative set of high-quality variants obtained using the same steps described in these methods from a subset of 39 high-quality samples. This method includes a variant calling approach that takes a consensus from GATK and bcftools and applies thresholds based on three variant quality annotations, selecting a conservative set of shared SNPs and Indels (FS<=2, MQ>=59 & QD>=20). The thresholds used were based on the distribution of these annotations across the sample set.

SNPs and indels were called using the consensus approach applied to outputs from GATK and bcftools variant callers using a modified version of a previously described workflow (76, 77). For GATK, HaplotypeCaller was used to identify potential variants in each sample, with the resulting GVCF files merged using CombineGVCFs, and join-genotyping performed using GATK’s GenotypeGVCFs. A similar joint-calling approach was implemented with bcftools using the mpileup and call subcommands. A consensus was taken of the resulting VCF files generating a conservative list of high-quality variants. Finally, SNPs and small indels were filtered using GATK’s VariantFiltration using the same thresholds outlined above.

### Data filtering

Further filtering was applied based on clonality, genotypic missingness and minor allele frequency (MAF), depending on the downstream analysis. For clonality, within-isolate fixation index F_WS_ (78) was calculated using the moimix package (github.com/bahlolab/moimix) and samples with F_WS_<0.85 were removed from downstream analyses (15). To better understand the relatedness of the clones within these mixed infections, the non-reference allele frequency (NRAF) was also plotted across the genome for individual samples using ggplot2 (79). Genotypic missingness and MAF were then calculated using PLINK2 (80, 81). SNPs with MAF <5% or genotypic missingness >25%, and samples with >25% genotypic missingness, were filtered from downstream analyses, as well as those located in hypervariable regions (Supplementary Table 2).

### Characterising population structure

To determine overarching population structure, several complementary strategies were employed including neighbour joining analysis based on identity by state (IBS), connectivity based on identity by decent (IBD), and ADMIXTURE analysis (82). IBS was calculated with PLINK and visualised with neighbour-joining trees (NJT) in R using ggplot2 and ggtree (83). IBD was calculated with hmmIBD (84), which implements a hidden Markov model to determine sequence segments of shared ancestry. Base R and igraph (85) were used for IBD visualisation at a variety of thresholds (represent the percentage of the genome that is IBD between pairs of samples). Thresholds for IBD connectivity plots were determined by halving 95% IBD and every subsequent threshold. To determine the proportions of mixed ancestry, ADMIXTURE was used to implement a maximum likelihood estimation, which was then visualised in R using ggplot2. CV error was calculated prior to ADMIXTURE analysis to identify the optimal K value. As K=3 was deemed optimal, exhibiting a low cross-validation error compared to other K values determined by ADMIXTURE’s cross-validation procedure, and the distribution of samples aligned with the NJT, the K clusters were referred to throughout the manuscript with the previously defined Peninsular- and macaque-associated cluster names (*Macaca fascicularis* (*Mf)*, *Macaca nemestrina* (*Mn)* & *Peninsular*).

### Identifying the presence of introgression events

We performed a bespoke analysis to identify possible genomic regions of introgression. First, we identified the major allele for each cluster at each genomic coordinate. Then using a sliding window approach (10kb windows), we determined the genetic distance for each sample to each cluster. Genetic distance is defined as the proportion of mismatched SNPs per sliding window (10kb) when comparing the called allele in the sample to the major allele for a cluster at each position. The genetic distances were then plotted on a two-dimensional axis, with different clusters along the x and y axis, and two-dimensional kernel density estimations (contours) were calculated for the genomic clusters using ggplot2 (github.com/tidyverse/ggplot2) and MASS (github.com/cran/MASS) packages, and the density contours overlayed on the plot. The *points.in.polygon* function (github.com/edzer/sp) was used to determine in which contours the windows are spatially located. Windows located within the contours of another cluster whilst outside the contours of their own were defined as introgressed. To be conservative, candidate windows underwent several filtering steps, including those that appear in >= 5% of the population and the removal of windows that overlap hypervariable regions (Supplementary Table 2).

### Exploring links between introgression and environmental land types

We performed subsequent regression analyses to explore whether surrounding village-level environmental land types and predicted vector habitat suitability are associated with *P. knowlesi* introgression. The primary residential addresses for deidentified *P. knowlesi* cases for the preceding 3 weeks before health facility presentation was first used to obtain centroid village-level location coordinates cross-checked for accuracy using Google Earth (version 7.3). Selected environmental classification metrics of forest fragmentation (percentage of Landscape – tree cover and Perimeter-Area Ratio – tree cover) within a 5km radius surrounding village locations were then calculated from a composite landscape metrics tool encompassing ESRI 2020 and Sentinel-2 GIS data at 10-metre resolution (86) (Supplementary Figures 2 & 3). The relative predicted *Anopheles* Leucosphyrus Complex mosquito vector occurrence surface from Moyes et al. (50) based on boosted regression tree models encompassing mosquito sampling presence/absence data and environmental covariates indicating habitat suitability was obtained through the *malariaAtlas* R package (87). The mosquito vector habitat suitability surface was averaged within a 5×5km grid around the geolocated village sites. Moran’s I was calculated to exclude spatial autocorrelation of the environmental land types and predicted vector habitat suitability at the selected grids. Univariate regression analyses were initially used to assess potential associations between these environmental parameters and the macaque-derived clusters. Tertiles were then generated representing the degree of introgression in samples, with samples categorised as having *low*, *medium*, or *high* introgression. Logistic regression models were subsequently implemented to assess if landscape fragmentation indices or *Anopheles* Leucosphyrus Complex habit suitability were associated with either the presence of the top ten most frequently introgressed windows (binomial) or the degree of introgression (ordinal). The Akaike Information Criterion (AIC) was compared to determine the optimal model design (scripts available in the attached GitHub repository).

### Identification of orthologous antimalarial drug resistance markers

Antimalarial drug resistance markers for P. falciparum and P. vivax were collated using multiple sources (33, 34) and their orthologues in P. knowlesi identified. This includes dihydrofolate reductase (*dhfr*), dihydropteroate synthase (*dhps*), chloroquine resistance transporter (*crt*), multidrug resistance protein 1 (*mdr1*), multidrug resistance-associated proteins 1 (*mrp1*), plasmepsin 4 (*pm4*), kelch 13 (*k13*), reticulocyte binding protein 1a (*rbp1a*) and reticulocyte binding protein 1b (*rbp1b*). *P. falciparum* and *P. vivax* orthologues were identified using PlasmoDB’s (88) *Orthologue and synteny* tool, which is based on the OrthoMCL database, a genome-scale algorithm for grouping orthologous protein sequences (89). Multiple sequence alignment was then performed on PlasmoDB sequences, comparing the *P. knowlesi* amino acid sequences against the relevant orthologues in *P. falciparum* and *P. vivax* to identify shared mutations between orthologue genes.

### Characterising subpopulations within Malaysian Borneo

Once the larger *P. knowlesi* population structure was characterised, we investigated differences between sub-population clusters within Malaysian Borneo and interrogated the genome of all relevant samples for signs of differentiation. Subpopulations were characterised by sub-setting the samples to downscale the major genomic clusters (*Mf* and *Mn*) of Malaysian Borneo, to control for population structure. Then, both IBS and IBD analyses were repeated to determine whether subpopulations exist within these larger populations. PLINK was used to calculate the fixation index (F_ST_), a measure of population differentiation due to genetic structure, specifically, the variance of allele frequencies between populations. ggplot2 was used to visualise F_ST_ across the genome using a non-overlapping sliding window (1-kb) approach. This allowed the identification of outlier regions, which were annotated to identify potential genes of interest. Further details on the methods described here, including the entire workflow and associated scripts can be found at https://github.com/JacobAFW/Pk_Malaysian_Population_Genetics.

## Financial Disclosure Statement

Sample collection and sample processing were supported by the Ministry of Health, Malaysia (grant number BP00500420 and grant number BP00500/117/1002); the Australian National Health and Medical Research Council (grant numbers 496600, 1037304 and 1045156); the US National Institutes of Health (grant numbers R01AI116472-03 and 1R01AI160457-01), and the UK Medical Research Council, Natural Environment Research Council, Economic and Social Research Council, and Biotechnology and Biosciences Research Council (grant number G1100796).

Whole genome sequencing was supported by a Singaporean Ministry of Education Grant (grant number MOE2019-T3-1-007), and salary support for bioinformatics and analyses through an Australian NHMRC Ideas Grant (grant number APP1188077).

MF and MG were supported by NHMRC Emerging Leader 2 fellowships; MG was also supported by the Australian Centre for International Agricultural Research and Indo-Pacific Centre for Health Security, Department of Foreign Affairs and Trade, Australian Government funded ZOOMAL project (LS/2019/116).

The funders had no role in study design, data collection and analysis, decision to publish, or preparation of the manuscript.

## Acknowledgements

We thank the study participants, and the research team at the Infectious Disease Society Kota Kinabalu Sabah including Sitti Saimah binti Sakam, Danshy Anne Alaza, Azielia Elastiqah binti Salamth and Mohd Rizan Osman. We thank the Director-General, Ministry of Health, Malaysia, for permission to publish this manuscript. We thank Dr Freya Shearer and Dr David Duncan from the University of Melbourne for their consultation on specific analyses.

## Ethics Approval Statement

The research was performed in accordance with the Declaration of Helsinki and ethics approval was obtained from the medical research ethics committees of the Ministry of Health, Malaysia and Menzies School of Health Research, Australia.

## Data Availability Statement

Genomic data produced as part of this work are available at the Sequence Read archive (SRA) of the National Center for Biotechnology Information (NCBI) under the BioProject ID PRJNA1066389 and all bioinformatic and analytical scripts are available at https://github.com/JacobAFW/Pk_Malaysian_Population_Genetics.

## Conflict of Interest Disclosure

None of the authors have any conflicts to disclose.

## Supporting Information

S1 Text. Masked genomic regions and supplementary outputs.

## Notes

### Competing Interest Statement

The authors have declared no competing interest.

